# Bull Trout passage at beaver dams in two Montana streams

**DOI:** 10.1101/2022.09.10.507435

**Authors:** J. Marshall Wolf, Niall G. Clancy, Leo R. Rosenthal

## Abstract

Beaver (*Castor canadensis*) translocation and mimicry is an increasingly popular tool for process-based restoration of degraded streams. Processes influenced by beaver restoration include stream-floodplain connectivity, multi-threaded channel formation, enhanced riparian condition and fire resiliency, and streamflow attenuation. Beaver activity can also be a nuisance in agricultural settings, increase invasive species abundance, and create in-stream barriers to fish. Previous studies indicate that spring-spawning salmonid species can pass beaver dams in higher proportions than fall-spawning species. Thus, restoration or mimicry of beavers in streams containing fall-spawning, threatened Bull Trout (*Salvelinus confluentus*) is of concern to many biologists. We evaluated Bull Trout passage at beaver dams in two Montana streams: Meadow Creek (East Fork Bitterroot River drainage) in summer 2020 and Morrison Creek (Middle Fork Flathead River drainage) from 1997 to 2011. In Meadow Creek, 16% of PIT-tagged Bull Trout which entered a large beaver dam complex were detected upstream of some dams, but no fish moved through the entire 1 km complex. In Morrison Creek, redds were more likely than random to be located below dams, but two or more redds were found upstream of at least one dam in 6 of 9 years dams were present. These results suggest that beaver dams can affect the movement of Bull Trout, but that passage depends on the characteristics of individual dams and reach geomorphology. Our methods cannot distinguish between inhibition of fish movement and selection of beaver-created habitats by fish. Therefore, we suggest future research on beaver restoration in streams with Bull trout.

## Introduction

Beaver (*Castor sp*.) dams have profound impacts on stream habitats (Westbrook et al. 2011; Levine and Meyer 2019) including increasing late-season water flow (Nyssen et al. 2011; Majerova et al. 2015), resiliency to wildfires (Fairfax and Whittle 2020), and habitat for native co-occurring species (Cook 1940; Rosell et al. 2005) while also interfering with infrastructure (Albert and Trimble 2000) and providing habitat that can be beneficial for non-native fishes in some locations (Gibson et al. 2015). Historical removal of beavers from across North America and a growing recognition of the benefits they provide has led to a dramatic upswing in the use of beaver restoration and mimicry to restore degraded ecosystems in recent years (McKinstry and Anderson 2002; Pilliod et al. 2018; Wheaton et al. 2019). Beaver dam analogs (BDAs) are being widely deployed to aggrade streams and improve habitat for beaver recolonization in systems where beaver extirpation has occurred (Pollock et al 2014; Pilliod et al. 2018). Most salmonids in North America and Europe co-evolved with beavers, and beaver dams have many direct, positive impacts on salmonids including creation of high-quality habitat for adults and juveniles (Cook 1940; White and Rahel 2008), providing thermal refuge during warm summer months (Weber et al. 2017) and anchor-ice-free winter refuges (Jakober et al. 1998; Lindstrom and Hubert 2004). However, beaver dams are known to impede salmonid movement under some circumstances (DuPont et al. 2007; Lokteff et al. 2013), and the extent to which this occurs, especially during spawning migration, is of particular interest to fish biologists (Kemp et al. 2012). If beaver dams impede salmonid movement during critical life-history periods, special consideration may be needed for streams with threatened salmonids for which short-term recruitment is just as critical as long-term habitat restoration.

A number of previous studies have examined salmonid passage at beaver dams. Native Bonneville Cutthroat Trout (*Oncorhynchus clarkii utah*) in northern Utah passed beaver dams more effectively than non-native Brown Trout (*Salmo trutta*) or Brook Trout (*Salvelinus fontinalis*) (Lokteff et al. 2013). Adult and juvenile steelhead (*O. mykiss*) were unhindered by beaver dams and BDAs in northeastern Oregon (Bouwes et al. 2016), and native Arctic Grayling (*Thymallus arcticus*) in southwestern Montana passed beaver dams successfully on 88% of their attempts (Cutting et al. 2018). However, Cutthroat Trout, steelhead, and Arctic Grayling all spawn in the spring when streamflow is near its yearly maximum in the Western US, and thus passage at channel-spanning structures is most probable. Indeed, studies of fall-spawning Atlantic Salmon (*Salmo salar*) have found occasional-to-frequent blockage of upstream movement by beaver dams in New Brunswick (Mitchell and Cunjack 2007) and Nova Scotia where passage varied predictably with total autumn precipitation (Taylor et al. 2010). Similarly, redd counts of Sea Trout (*S. trutta trutta*) in Lithuania were greater in 500 m reaches below large beaver dams than above, whereas streams with only smaller River Trout (*S. trutta fario*) redds were distributed almost evenly above-and-below smaller, less-intact dams (Kesminas et al. 2013). The passage of juvenile Coho Salmon (*O. kisutch*) at BDAs in northern California was found to vary seasonally with streamflow levels (O’Keefe 2021). As such, there is still question as to whether beaver dams or BDAs could act as barriers to species that spawn during the lower limb of the hydrograph.

Bull Trout (*Salvelinus confluentus*) are coldwater specialists and fall spawners that have been listed as a threatened species under the Endangered Species Act in the United States since 1998. Primary causes of decline are habitat degradation, invasive species, warming water temperatures, and stream fragmentation due to dams, diversions and dewatering (Rieman and McIntyre 1993; Nelson et al. 2002; Al-Chokhachy et al. 2016). Bull Trout abundances have substantially declined throughout their range and, in western Montana, most river basins that once contained large-bodied migratory individuals now contain only isolated, headwater-resident populations (MBTSG 1995). Only the upper Flathead and upper Kootenai River basins still contain relatively abundant populations of migratory Bull Trout (Kovach et al. 2018). Due to these range-wide declines, further fragmentation of Bull Trout populations by channel spanning structures such as beaver dams is of great concern. To help determine if process-based restoration utilizing beaver translation or mimicry is viable for usage in streams with Bull Trout, we examined Bull Trout passage at natural beaver dams in tributaries to the East Fork Bitterroot River and Middle Fork Flathead River.

## Methods

### Study Area

Meadow Creek is a third-order tributary of the East Fork Bitterroot River in the Sapphire Mountains of western Montana (Figure 1) with an estimated average baseflow at the project area near 0.1 m^3^/s with average peak flows estimated at about 0.71 m^3^/s (McCarthy et al. 2016). A large beaver dam complex approximately 1 km long is located at stream km 6.4. Morrison Creek is a third-order tributary to the Middle Fork Flathead River (Figure 1) with an estimated average baseflow of 0.71 m^3^/s and average peak flow greater than 8.5 m^3^/s (McCarthy et al. 2016). Both contain healthy Bull Trout populations.

**Figure 1.**
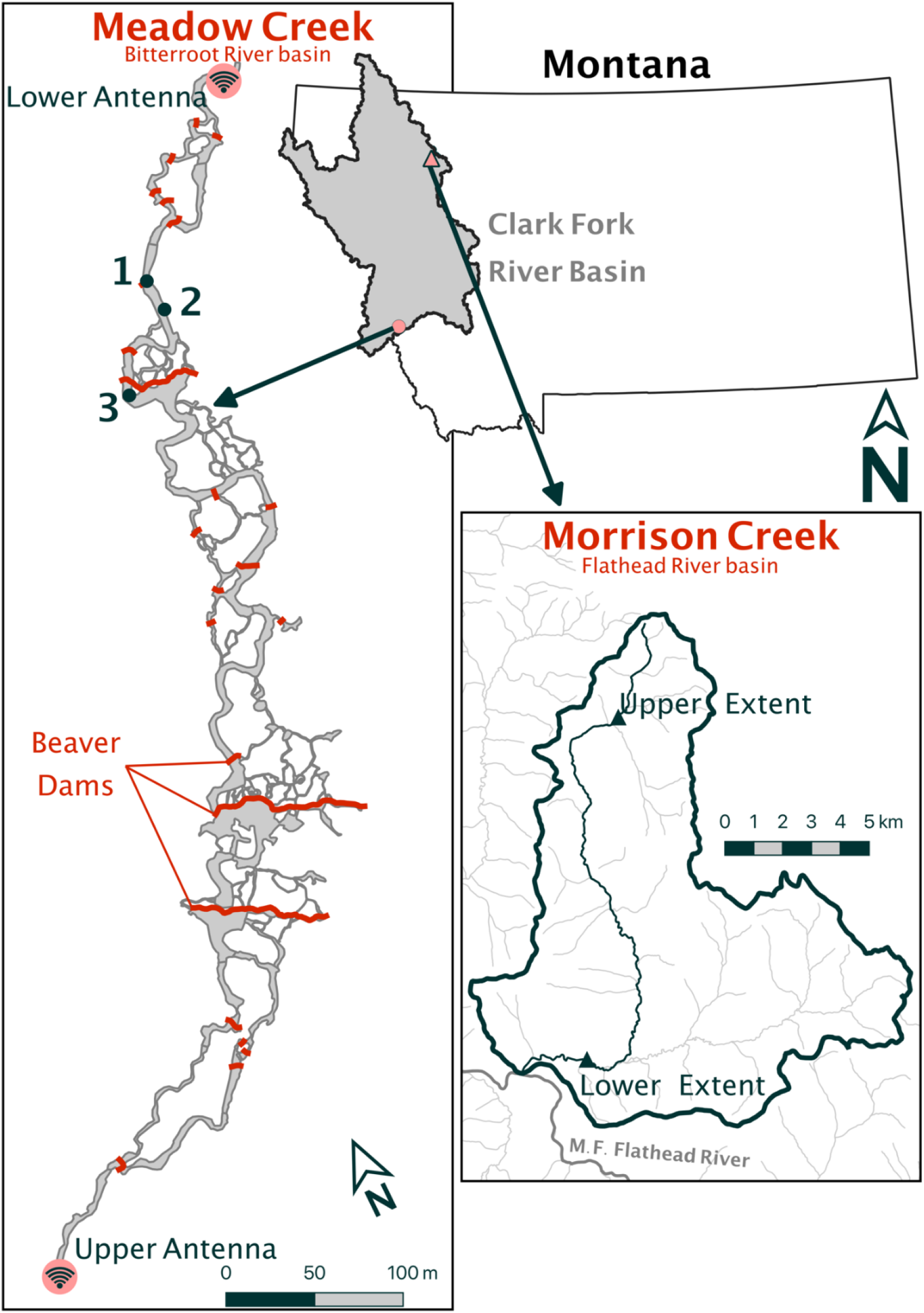
Study sites within the Clark Fork River basin. The left side displays locations of the beaver dams, PIT tag antennas, and fish detected with the mobile PIT tag reader (numbers 1-3 correspond to fish descriptions in results section) at Meadow Creek, East Fork Bitterroot River, MT. Right side displays the redd survey extents within Morrison Creek, Middle Fork Flathead River, MT

### Meadow Creek PIT Tag Study

On July 21st, 2020, we installed a battery-powered, submersible passive integrated transponder (PIT) tag antenna system (Biomark©, Boise, ID) on the up- and downstream ends of the Meadow Creek beaver dam complex (Figure 1). We placed rocks around the antennas to force PIT-tagged trout to swim close enough to the antenna to be recorded. In the three days following installation, we captured 49 Bull Trout via backpack electrofishing (LR-24 Backpack Shocker, SmithRoot©, Vancouver, WA) in and above the beaver dam complex. We implanted 37 individuals larger than 100 mm with a 12 mm, uniquely-coded PIT tag (Biomark© Model HDX12) and released fish immediately below the downstream antenna. Antenna batteries were replaced bi-weekly to ensure continuous antenna operation until they were removed on October 1^st^. We chose start and end dates based on a telemetry study in nearby Skalkaho Creek that found the vast majority of upstream Bull Trout spawning runs occurred between late July and September (Clancy 2017).

At study completion, we surveyed the entire beaver dam complex with a mobile PIT tag antenna (Biomark© BP Plus Portable Antenna) which recorded the location and tag number of all detected fish as we waded through the water in a similar manner to electrofishing. At that time, we also censused beaver dams in the complex by recording the locations, condition, crest heights, and jump heights following the Hafen et al. (2020) beaver dam rapid assessment method. We calculated a pool depth-to-jump height ratio (e.g. ratio = pool depth / jump height; hereafter pool-to-jump ratio) for each dam (Stuart 1964; Kondratieff and Myrick 2006). Aerial imagery was acquired with a drone (DJI, Shenzhen, China) and we used this imagery to delineate dam crests in ArcGIS Pro (ESRI, Redlands CA, USA).

### Morrison Creek Redd Counts

Standardized Bull Trout redd counts have occurred in the Middle Fork Flathead River tributaries since 1979 and serve as an index of Bull Trout abundance in the watershed (Shepard and Graham 1982). In most years, trained redd surveyors walk Morrison Creek (typically upstream to down) from approximately stream kilometer 16.9 to 3.1 (13.8 km total). During “basin-wide survey” years, a longer reach, from stream kilometer 20.1 to 3.1 (17.0 km total), is surveyed. Using archived 1997 to 2011 field notes, we converted recorded pace counts of each redd and beaver dam location to approximate stream kilometer; we used year-specific pace-to-kilometer conversions by dividing total paces in a given year by the known kilometers surveyed. Eleven years were able to be converted in this way. Beavers were trapped out of Morrison Creek in 2010 due to concerns regarding trout passage and several dams were breached in the study area.

To determine if redd locations were randomly distributed below beaver dams, we randomly drew the same number of locations as redds present for that year from a number line representing 1-meter stream increments of the survey reach. For both real redds and randomly selected locations, we calculated the distance downstream from a beaver dam, which were negative if upstream of all dams. We then compared the distribution of real redds and random locations across all years using two Kolmogorov-Smirnov tests to determine if the real redd locations were randomly distributed. The first test was for random distribution with respect to distance downstream from a dam. The second test was for random distribution with respect to stream meters along our survey reach. We ran two sided Kolmogorov-Smirnov tests with α set at 0.05 to test the null hypothesis that both samples were drawn from the same distribution (Conover 1972). All analyses were conducted in the program R (R Core Team, 2022).

## Results

### Meadow Creek PIT Tag Study

The mean length of the 37 tagged fish was 155.5±49.3 mm, and 6 fish were larger than 200 mm (Figure 2). We detected 18 (49%) of the Bull Trout at the downstream antenna (Figure 2). Within eight days of tagging (by July 29^th^, 2020), 9 fish had passed the lower PIT tag antenna. The remaining 9 fish were detected at a relatively steady rate over the next 3 weeks with the final new detection occurring on August 13^th^. No fish were detected at the lower antenna between September 14^th^ and when it was removed on Oct 1^st^. Of the six 200+ mm fish tagged, four (66.67%) were detected at the lower antenna while 14 of the 31 (45.2%) fish under 200 mm were detected (Figure 2). No fish were detected at the upstream antenna.

**Figure 2.**
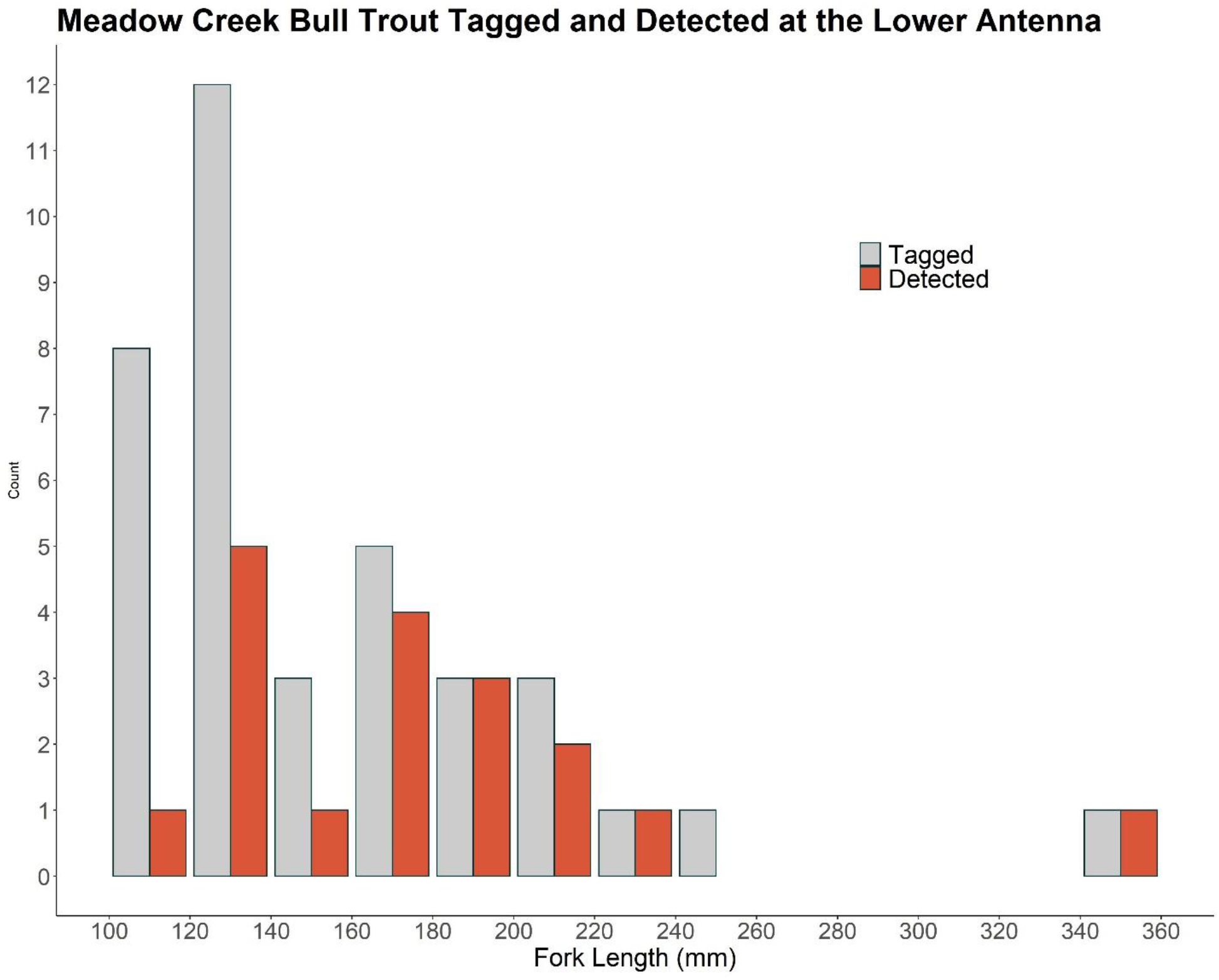
Length distribution of Bull Trout caught, PIT tagged, and released below the lower antenna in grey along with those detected by the lower antenna in red

Three fish with lengths of 117, 107, and 131 mm were detected with a mobile PIT antenna on October 1^st^. Two of these fish were detected above several secondary dams (max jump height 0.5 m, Figure 1: fish #1 and #2) and the other fish was detected just above the first primary dam (jump height 0.5m, Figure 1: fish # 3). The mean pool-to-jump ratio of these dams was 1.40±0.91m (range 0.5 - 2.5m, Table S1).

We recorded 35 beaver dams along the 1 km study reach of Meadow Creek (Table S1). Of these dams, 3 (9%) were primary dams creating a pond with a beaver lodge or food cache, 27 (77%) were intact secondary dams, and 5 (14%) were blown out or breached. Crest and jump heights of primary dams were 1.2±0.17 m (mean±standard deviation) and 0.82±0.30 m, respectively. Secondary dams had crest heights of 0.74±0.28m with jump heights of 0.38±0.27 m. The maximum observed jump height was 1.1 m. Along the lateral margins of all three primary dams, we identified side channels with lower jump heights than the primary dam crest, which may have served as alternative routes for upstream fish movement (Figure 1, see side channels).

### Morrison Creek Redd Counts

Of the 11 years we analyzed, only in 1998 and 2011 (the year after beavers were trapped) were no beaver dams recorded within the Morrison Creek study area (Figure 3). During this time period, the monitoring reach averaged 28.6±12.8 redds. In every year except 1998, the majority of Bull Trout redds were located in the lower half of the study area, below stream kilometer 9.3. Every year with beaver dams had multiple redds requiring fish to pass two or more dams, except for 2003 and 2004. Those years had a single redd that would require multiple dam passage and the lowest redd counts of the dataset (10 and 14 respectively; Figure 3 and Table 1). Redds were found above some beaver dams during all 9 years they were present. In 6 of the 9 years with beaver dams, the majority of redds were downstream of all beaver dams (i.e. not above any dam; Table 1). No redds were recorded above the upstream most dam in 7 of the 9 years with dams (Table 1, Figure 3). Across all years, while an average of 14% of the survey reach was half-a-kilometer or less below a beaver dam, 39% of redds were half-a-kilometer or less below a dam._A Kolmogorov-Smirnov (K-S) test on the distance downstream from a dam concluded that observed redds were not drawn from the same distribution as the random locations (D = 0.15972, p-value = 0.001289). A second K-S test comparing stream meter locations revealed that the observed redds were not randomly located within the stream (D = 0.28035, p-value = 3.097e-12).

**Figure 3.**
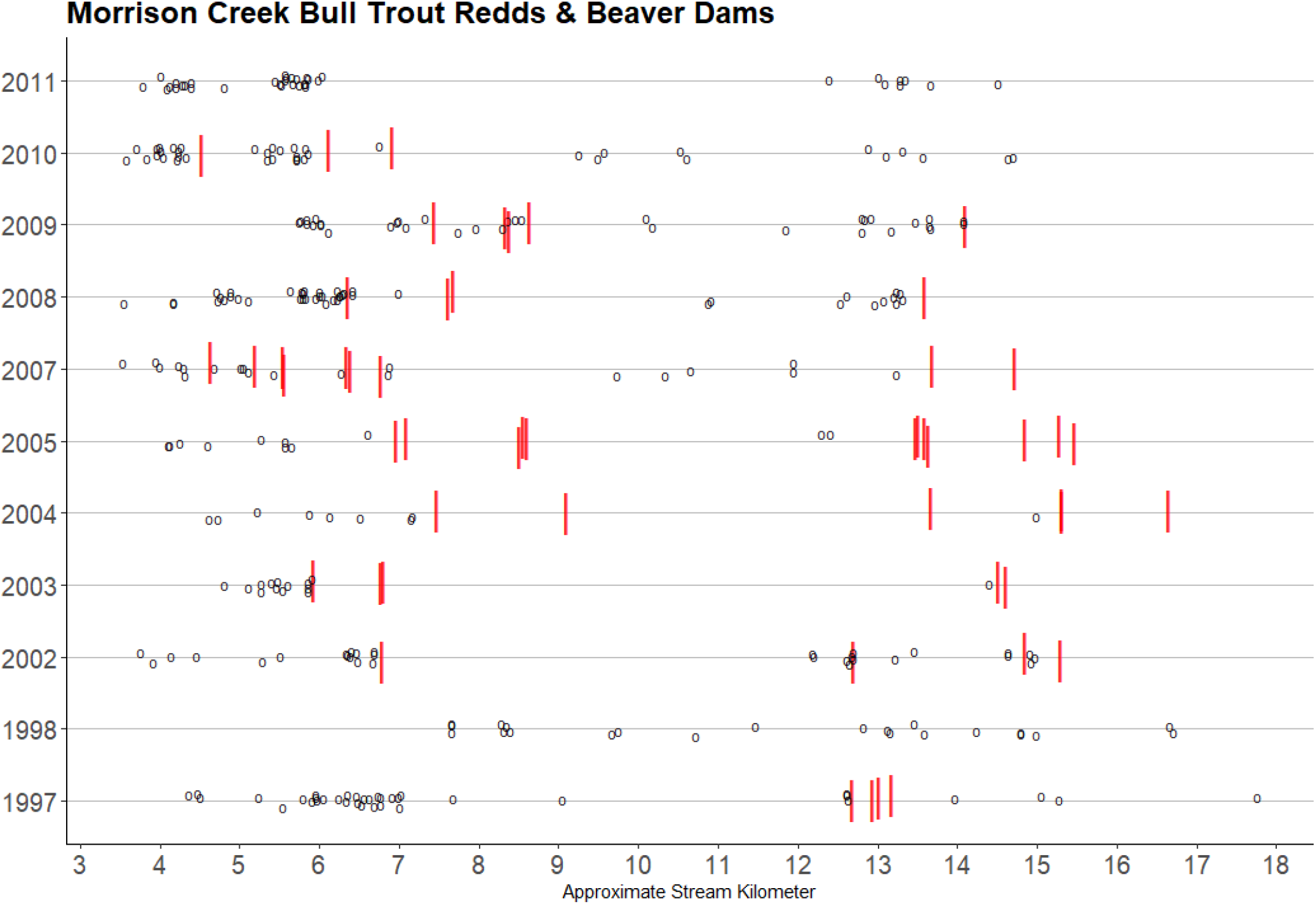
Morrison Creek redd counts (downstream left), Flathead River basin, Montana, 1997 - 2011. Years included are those in which beaver dams were recorded and pace counts could be converted to stream kilometers. Red lines indicate locations of beaver dams and circles indicate Bull Trout redds. Kilometers proceed upstream. Both redds and dams are offset on the Y axis by a small amount to enhance visibility of features in close proximity.

**Table 1.** Location of Bull Trout redds in proximity to beaver dams during each of the 1997-2011 redd count surveys in Morrison Creek, Middle Fork Flathead River basin. Numbers in parentheses refer to yearly proportion. “Immediately below dam” refers to 0.5 kilometers below beaver dams. Above KM 9.3 (stream kilometer 9.3) is provided as an index of the spread of redds. *Beaver dams were recorded both individually and as a complex and the number of dams each year should also be viewed as an index.

| Year | Beaver<br>Dams* | Reach proportion<br>immediately below dam | Number of Redds |  |  |  | Total |
| --- | --- | --- | --- | --- | --- | --- | --- |
|  |  |  | Immediately<br>below dam | Above all<br>dams | Below all<br>dams | Above<br>KM 9.3 |  |
| 2011 | 0 | 0 | 0 | No Dams | No Dams | 9 (0.23) | 39 |
| 2010 | 3 | 0.11 | 16 (0.37) | 12 (0.28) | 17 (0.40) | 11 (0.26) | 43 |
| 2009 | 5 | 0.13 | 13 (0.38) | 0 | 14 (0.41) | 14 (0.41) | 34 |
| 2008 | 4 | 0.11 | 18 (0.39) | 0 | 31 (0.67) | 12 (0.26) | 46 |
| 2007 | 9 | 0.27 | 11 (0.52) | 0 | 7 (0.33) | 6 (0.29) | 21 |
| 2005 | 12 | 0.22 | 1 (0.07) | 0 | 13 (0.87) | 2 (0.13) | 15 |
| 2004 | 6 | 0.18 | 4 (0.40) | 0 | 9 (0.90) | 1 (0.10) | 10 |
| 2003 | 5 | 0.10 | 9 (0.64) | 0 | 13 (0.93) | 1 (0.07) | 14 |
| 2002 | 4 | 0.14 | 19 (0.61) | 0 | 17 (0.55) | 14 (0.45) | 31 |
| 1998 | 0 | 0 | 0 | No Dams | No Dams | 16 (0.70) | 23 |
| 1997 | 4 | 0.06 | 4 (0.10) | 4 (0.1) | 36 (0.9) | 8 (0.20) | 40 |

## Discussion

Given that no Bull Trout passed all dams in Meadow Creek and redd locations in Morrison Creek were more likely than random to be immediately downstream of dams, we surmise that beaver dams affect the movement and spawning locations of Bull Trout. We are unable to determine if this is a result of impairment of upstream migration or selection of downstream habitats. For example, during 2011 after beaver dams were breached in Morrison Creek, Bull Trout still spawned at high densities within the same reaches that were previously below beaver dams even though they were newly passable (2011, Figure 3). Differentiating between blockage and selection is further complicated by the fact that Bull Trout prefer redd locations with active hyporheic exchange (Baxter and Hauer 2000), and beaver dam complexes (Westbrook et al. 2006, Weber et al. 2017) increase the magnitude and extent groundwater-surface water interactions.

Our results are similar to findings of a report from northern British Columbia that found Bull Trout redds both above and below beaver dams up to 1.5 m in height on three different creeks. Passage at those dams, like at the ones reported here, appeared to vary yearly and seasonally based on flow conditions and dam morphology (Bustard 2017). Only a well-designed experiment in combination with laboratory studies could determine if Bull Trout are impaired from upstream migration or selecting downstream habitats created by dams in the form of pools, hyporheic flow paths, or spawning gravels. It is likely that both impairment and selection occur simultaneously, similar to the manner in which beaver dams are known to both create suitable rearing habitat for Bull Trout (Jakober et al. 1998) and are thought to delay or halt downstream migrations in low-flow conditions (DuPont et al. 2007).

Laboratory studies found that a pool-to-jump ratio of approximately 0.8 is ideal for salmon and trout (Stuart 1964). Our case study generally corroborates this number for Bull Trout, with passage detected at dams with pool-to-jump ratios between 0.5 and 2.5 (Table S1). Specific estimates of optimal pool-to-jump ratios or maximum jump heights of Bull Trout have not been calculated, to our knowledge. Two primary dams along the thalweg route did not have any passage detected. Their observed pool-to-jump ratios were 0.27 and 0.29, much lower than other dams we observed passage of or the 0.8 optimal estimate indicating that they would likely be difficult for Bull Trout to pass. Side channels along both of these primary dams provided smaller jump heights (minimum: 0.23 and 0.15) but similar pool-to-jump ratios (0.20 and 0.35). Studies of closely related Brook Trout jumping mechanics indicate pool-to-jump ratios are much less important for fish larger than 200 mm attempting short jumps of < 20 cm (Kondratieff and Myrick 2006). Several studies have also observed salmonids swimming directly through or around (via small rivulets) beaver dams (Lokteff et al. 2013; Cutting et al. 2018). It is likely that Bull Trout in both Meadow and Morrison Creek would have had to go over or through some beaver dams.

The exact timing of Bull Trout migration and spawning varies by location and year (Rieman and McIntyre 1993, 1995; Swanberg 1997). Bull Trout in the East Fork Bitterroot River typically spawn in September (Jakober et al. 1998; Nyce 2011), but have been observed spawning as early as the end of August (M. Jakober, U.S. Forest Service, *personal communication*). Within Meadow Creek, a tributary to the East Fork, 80% of the Bull Trout detections at the lower antenna occurred prior to August 17^th^, a pattern similar to fish in nearby Skalkaho Creek (Clancy 2017). Bull Trout in other Montana rivers have been observed making spawning movements from May through September, with long-distance migratory fish moving earlier than resident fish (Fraley and Shepard 1989; Swanberg 1997; Paragamian and Walters 2011). Differences in movement timing, life histories, annual stream hydrology and floodplain connectivity (e.g. overflow and side channels) could lead to varying beaver dam passage rates for Bull Trout across different individual dams, systems, and years.

Further research on Bull Trout passage at beaver dams is necessary. It is possible placement of the upstream PIT tag antenna closer to the halfway point of the Meadow Creek beaver complex would be better for detecting fish that had passed lower dams but remained within the complex for spawning, overwintering or both. We have several lines of evidence for spawning activities taking place with the beaver complex. Yearly redd counts conducted within our study reach by the U.S. Forest Service indicate some passage of beaver dams by migratory Bull Trout in recent years (M. Jakober, U.S. Forest Service, unpublished data). During electrofishing, we captured one potentially fluvial fish (migrant from the East Fork Bitterroot) 347 mm total length from within the beaver complex. During our mobile PIT tag detection efforts in October 2020, we observed several large bodied Bull Trout in the 300-400mm range behind the two upstream most primary dams that were not observed during our initial electrofishing in July 2020. However, the complexity of multiple channel braids within the beaver dam complex and our limited number of PIT tag antennas prevented us from using a mid-complex antenna approach. Detection of fish within the complex was likely underreported by the mobile PIT tag antenna because it is not ideal for use in streams as large and deep as beaver-influenced Meadow Creek. Future studies using this method could consider combining electrofishing with mobile PIT tag reading to enhance recapture probabilities.

Based on the two case-studies we highlight in tributaries to the East Fork Bitterroot and Middle Fork Flathead Rivers, we suggest beaver restoration and mimicry within Bull Trout streams be implemented in an adaptive framework with frequent monitoring. Restoration efforts should endeavor to reconnect the stream with its floodplain (e.g. be built so as to create flooding or side channels around the dam), not only to maximize the restoration benefits (Burchsted et al. 2010; Pollock et al. 2014), but to also create multiple passage routes for fish species (Lokteff et al. 2013; Bouwes et al. 2016; Cutting et al. 2018). If maximizing the stream length available to spawning Bull trout is the top management concern, we recommend seasonal notching of beaver dams that have jump heights of greater than 0.6, pool-to-jump ratios of less than 0.8, and no side channel routes (Bustard 2017). This will promote increased passage (Taylor et al. 2010) while maintaining benefits for juvenile growth and development during the following spring and summer (White and Rahel 2008; Bouwes et al. 2016).

## Acknowledgements

We are indebted to Peter Mackinnon for providing needed equipment without which this project would not have been possible. Chris Clancy and Tara Gallagher were incredibly generous with their time, swapping antenna batteries throughout the project’s duration. Andrea Price and Grace Pierstorff of the Bitter Root Water Forum helped capture fish and attend to batteries, and Scott Hawxhurst provided valuable field support in the Flathead basin. Annual Bull Trout redd counts in the Flathead River basin have been a joint effort by numerous FWP personnel throughout the years. Thanks also to Mike Jakober and Jason Lindstrom for their advice and permitting help. N. Clancy was affiliated with MFWP at the time of fish sampling.

