## Supplementary material for "Bull Trout passage at beaver dams in two Montana streams": Figure S1 and Table S1

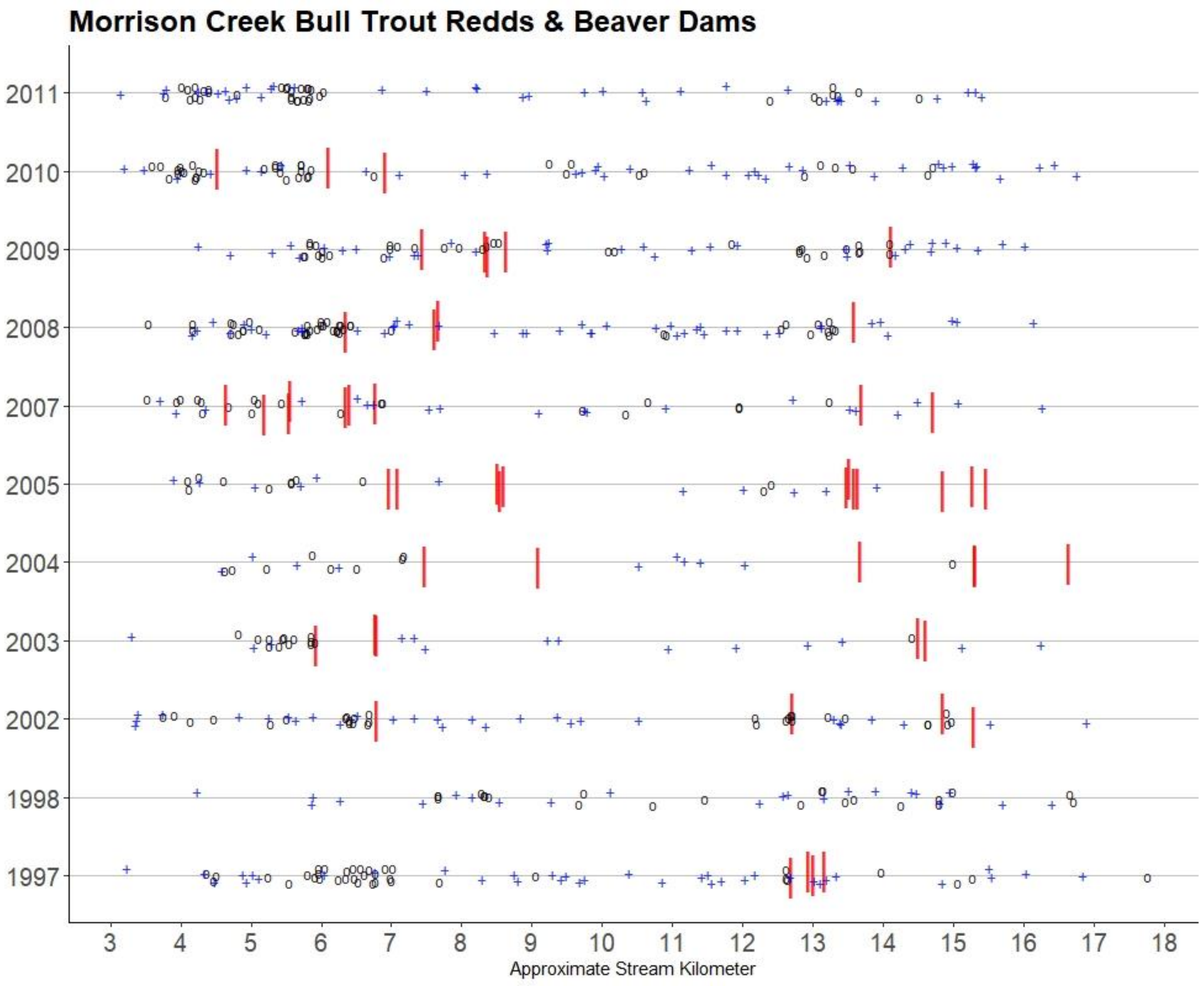


**Figure S1.** Morrison Creek redd counts (downstream left), Flathead River basin, Montana, 1997 – 2011 with randomly generated points (blue crosses) for comparison to actual redd locations (circles). Red lines indicate locations of beaver dams and circles indicate Bull Trout redds. Years included are those in which beaver dams were recorded and pace counts could be converted to stream kilometers Kilometers proceed upstream. Both redds and dams are offset on the Y axis by a small amount to enhance visibility of features in close proximity.

| Dam Number | Dam Type | Crest Height | Jump Height | P-J Ratio | Crest Length | Condition |
|---|---|---|---|---|---|---|
| 1 | secondary | 0.6 | 0.4 | 0.50 | 3.1 | intact |
| 2 | secondary | 1.45 | 1.3 | 0.50 | 3.2 | intact |
| 3 | secondary | 0.75 | 0.4 | 0.88 | 3.8 | intact |
| 4 | secondary | 0.95 | 0.25 | 2.80 | 2.25 | intact |
| 5 | secondary | 0.9 | 0.5 | 0.80 | 4.0 | intact |
| 6 | secondary | 0.8 | 0.4 | 1.00 | 6 | breached |
| 7 | secondary | 0.7 | 0.2 | 2.50 | 5 | breached |
| 8 | secondary | 0.85 | 0.25 | 2.40 | 5 | intact |
| 9 | secondary | 0.3 | 0.3 | 0.00 | 2.25 | blown out |
| 10 | primary | 1.1 | 0.5 | 1.20 | 31.5 | intact |
| 11 | secondary | n/a | n/a | n/a | n/a | blown out |
| 12 | secondary | 1.1 | 0.55 | 1.00 | 5 | intact |
| 13 | secondary | 0.6 | 0.4 | 0.50 | 3.3 | intact |
| 14 | secondary | 0.75 | 0.63 | 0.19 | 3.35 | intact |
| 15 | secondary | 0.8 | 0.5 | 0.60 | 12 | intact |
| 16 | secondary | n/a | n/a | n/a | n/a | blown out |
| 17 | secondary | 0.95 | 0 | n/a | 1.25 | intact |
| 18 | secondary | n/a | n/a | n/a | n/a | blown out |
| 19 | secondary | 1.05 | 0.65 | 0.62 | 2.4 | intact |
| 20 | secondary | 0.73 | 0.5 | 0.46 | 1.3 | intact |
| 21 | secondary | n/a | n/a | n/a | n/a | blown out |
| 22 | secondary | 0.71 | 0.55 | 0.29 | 1.95 | intact |
| 23 | secondary | 0.71 | 0 | n/a | 1.15 | intact |
| 24 | secondary | 0.45 | 0 | n/a | 0.6 | intact |
| 25 | secondary | 0.45 | 0 | n/a | 1.1 | intact |
| 26 | secondary | 0.31 | 0.25 | 0.24 | 0.3 | intact |
| 27 | secondary | 0.6 | 0.35 | 0.71 | .5 | intact |
| 28 | secondary | 1.4 | 0.75 | 0.87 | 10.6 | intact |
| 29 | primary | 1.4 | 1.1 | 0.27 | 89 | intact |
| 30 | primary | 1.1 | 0.85 | 0.29 | 87 | intact |
| 31 | secondary | 0.5 | 0.3 | 0.67 | 6 | intact |
| 32 | secondary | 0.4 | 0.24 | 0.67 | 5 | intact |
| 33 | secondary | 0.55 | 0.37 | 0.49 | 1.9 | intact |
| 34 | secondary | 0.75 | 0.4 | 0.88 | 0.45 | intact |
| 35 | secondary | 0.5 | 0.12 | 3.17 | 7.9 | breached |

**Table S1.** Survey of all beaver dams in Meadow Creek in summer 2020. Heights and lengths are in meters. Jump height is the crest height above the downstream pool. P-J ratio is the ratio of the pool depth to the jump height.
